# Variable optical properties of light-harvesting complex II revisited

**DOI:** 10.1101/2020.10.05.312405

**Authors:** Masakazu Iwai, Jie-Jie Chen, Soomin Park, Yusuke Yoneda, Eva M. Schmid, Daniel A. Fletcher, Graham R. Fleming, Krishna K. Niyogi

**Affiliations:** Molecular Biophysics and Integrated Bioimaging Division, Lawrence Berkeley National Laboratory, Berkeley, CA 94720, USA; Department of Plant and Microbial Biology, University of California, Berkeley, CA 94720, USA; Department of Chemistry, University of California, Berkeley, CA 94720, USA; Department of Chemistry, University of Science & Technology of China, Hefei, 230026, China; Department of Bioengineering, University of California, Berkeley, CA 94720, USA; Biological Systems and Engineering Division, Lawrence Berkeley National Laboratory, Berkeley, CA 94720, USA; Kavli Energy Nanoscience Institute, Berkeley, CA 94720, USA; Howard Hughes Medical Institute, University of California, Berkeley, CA 94720, USA

## Abstract

Understanding photosynthetic light harvesting requires knowledge of the molecular mechanisms that dissipate excess energy in thylakoids. However, it remains unclear how the physical environment of light-harvesting complex II (LHCII) influences the process of chlorophyll de-excitation. Here, we demonstrate that protein-protein interactions between LHCIIs affect the optical properties of LHCII and thus influence the total energy budget. Aggregation of LHCII in the dark altered its absorption properties, independent of the amount of prior light exposure. We also revisited the triplet excited state involved in light-induced fluorescence quenching and found another relaxation pathway involving emission in the green region, which might be related to triplet excited energy transfer to neighboring carotenoids and annihilation processes that result in photoluminescence. LHCII-containing liposomes with different protein densities exhibited altered fluorescence and scattering properties. Our results suggest that macromolecular reorganization affects overall optical properties, which need to be addressed to compare the level of energy dissipation.

## Introduction

Major light-harvesting complex proteins of photosystem II (LHCIIs) form a trimeric structure in which chlorophyll (Chl) and carotenoid (Car) molecules (*i.e*., lutein, neoxanthin, and violaxanthin) are bound in a specific arrangement^1^. Photoexcitation of Chl generates the singlet excited state (^1^Chl*), followed by ultrafast excitation energy transfer (EET) through neighboring Chls within LHCII^2^. The excitation energy of ^1^Chl* is eventually trapped at a photosystem reaction center (RC) where charge separation is induced and photosynthetic electron transport is activated. Alternatively, ^1^Chl* de-excitation can take place via emission of energy as fluorescence. Under high light (HL) conditions, saturation of photosynthetic electron transport generates a situation in which open RCs are not locally available, which results in an increased accumulation of ^1^Chl*. This situation considerably increases the yield of intersystem crossing from ^1^Chl* to the triplet excited state (^3^Chl*)^3^, which can then transfer the excited energy to a triplet ground state oxygen to produce singlet oxygen, a highly reactive species capable of photooxidative damage. To minimize ^3^Chl* formation, LHCII proteins de-excite ^1^Chl* by non-radiative relaxation mechanisms that are dependent on the interaction between neighboring pigments^4-8^. There are two different techniques generally used to investigate these diverse ^1^Chl* de-excitation mechanisms: i) ultrafast spectroscopy to observe the transient kinetics of EET among pigments on the order of picoseconds to nanoseconds; and ii) modulation fluorometry to observe the induction kinetics of Chl fluorescence quenching on the order of minutes to hours. The ^1^Chl* de-excitation by these non-radiative processes, known as non-photochemical quenching (NPQ), has long been studied to understand photoprotection mechanisms in photosynthetic organisms under HL conditions^9,10^.

Although countless efforts have improved understanding of molecular mechanisms of NPQ in LHCIIs, it is still uncertain whether or not Chl de-excitation mechanisms are activated differently due to the physiological landscape of LHCII. To understand the contributions of different NPQ mechanisms, it is necessary to consider how and which quenchers are activated in different physical conditions that occur under different amounts of light exposure. Solubilization of LHCII in a detergent micelle, aggregation without a detergent, and reconstitution in lipid membrane environments are ideal methods to systematically investigate such effects. In this study, we explored how different physical conditions of LHCII affect its optical properties and influence the processes of ^1^Chl* and ^3^Chl* de-excitation. Most of our experimental approach was based on observations of steady-state fluorescence kinetics under different amounts of light exposure *in vitro*. Through this study, we found another possible radiative process occurring in the green region, which shows a correlation with the quenching of Chl fluorescence induced by excess excitation energy. We discuss the possible situations that can be compared between *in vitro* and *in vivo* conditions, which may illuminate complications associated with understanding the different components of NPQ and reconcile the various mechanisms proposed in the literature.

### The optical properties of LHCII are changed upon aggregation

We first used solubilized trimeric LHCIIs isolated from spinach thylakoid membranes to observe the kinetics of Chl fluorescence yields in the dark. As expected, the fluorescence yield stayed constant because LHCII remained solubilized in detergent micelles during the modulation fluorometry measurement (**Fig. 1a**). The unchanged fluorescence yield also indicates that the measuring light (ML) was weak enough to cause no further accumulation of ^1^Chl*. When we diluted the detergent concentration by putting the solubilized LHCII into the measuring buffer without the detergent, the fluorescence intensity gradually decreased (**Fig. 1a**), which reflects a typical process of LHCII aggregation^11^. We confirmed that the intensity of ML did not influence the extent of the decreased fluorescence yield (**Fig. 1b**), suggesting that the gradual decrease of the fluorescence intensity is caused exclusively by the process of aggregation. Adding the detergent back to the buffer caused a rapid increase of the fluorescence intensity to the same level as observed in the solubilized LHCII (**Fig. 1a**), indicating that the process of aggregation causes no prolonged modification of intrinsic properties of LHCII. When we removed the detergent completely from LHCII by dialysis, we observed much lower fluorescence intensity due to the highest degree of LHCII aggregation (**Fig. 1a**). Also, no change in the fluorescence yield was observed in the dialyzed LHCII, indicating that there was neither further aggregation nor change in intrinsic properties generated in the dark. These results suggest that the fluorescence intensity decreases due to the physical process of LHCII aggregation independent of environmental light conditions.

**Fig. 1.**
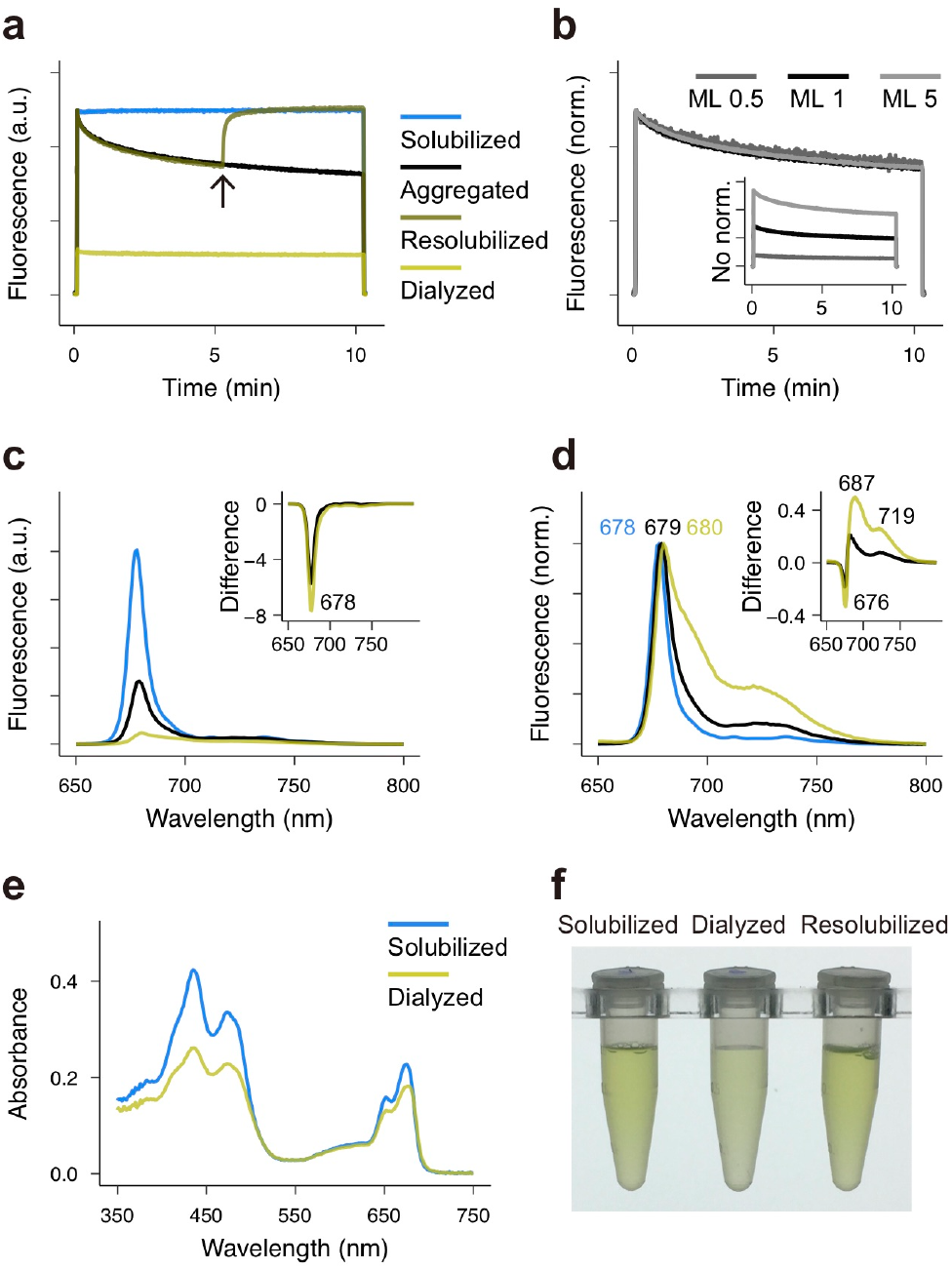
Aggregation-induced fluorescence decline in LHCII in the dark. **a**, Representative kinetics of Chl fluorescence yields observed using different LHCII samples in the dark. The solubilized LHCII stayed solubilized in the buffer with α-DDM at 0.03% (w/v), which is above the critical micelle concentration (CMC). The aggregated LHCII was gradually formed by putting the solubilized LHCII in the buffer without α-DDM, whose final concentration became 0.0004% (w/v), which is below the CMC. The resolubilized LHCII was formed from the aggregated LHCII by adding 0.03% (w/v) α-DDM to the buffer at the point indicated with an arrow. The dialyzed LHCII was prepared by removing α-DDM completely from the solubilized LHCII by dialysis (see Methods for details and Supplementary Fig. 1b). Chl concentration of each LHCII sample was adjusted to 5 µg Chl/mL. The intensity of ML was ∼1 µmol photons m^−2^ s^−1^. **b**, Chl fluorescence yields observed during aggregation measured using different intensities of ML. The number indicates the intensity in µmol photons m^−2^ s^−1^. The kinetics are normalized at the highest yield of each. The inset shows the kinetics without normalization. **c**,**d**, Representative Chl fluorescence spectra measured at 77 K normalized at the fluorescence peak (530 nm) of Alexa Fluor 430 (2 nM) added to the sample as an internal standard (**c**) or the main Qy peak (**d**). Excitation wavelength was 440 nm. The insets show the difference spectra from the spectra of the solubilized LHCII. Line colors are the same as in (**a**). **e**, Absorbance spectra measured using the solubilized LHCII and the dialyzed LHCII, adjusted to equal Chl concentration quantified by extraction in acetone. **f**, Photograph showing solubilized LHCII, dialyzed LHCII, and the same dialyzed LHCII resolubilized with 0.03% (w/v) α-DDM. Chl concentration of each sample was adjusted to 5 µg Chl/mL.

Using the same LHCII samples, we measured the steady-state Chl fluorescence spectra at 77 K. We examined the results using two different ways of normalization: i) normalized by the emission at 530 nm from an internal standard (Alexa Fluor 430) to observe changes in fluorescence intensities of overall spectra and ii) at the main Chl fluorescence peak (Qy band) to observe changes in spectral shapes. When the fluorescence spectra were normalized by the internal standard, the spectra of the aggregated and dialyzed LHCII samples exhibited a large decline at the Qy band compared to the solubilized LHCII (**Fig. 1c**). When the spectra were normalized at the Qy band, a shoulder of red-shifted emission appeared at ca. 720 nm in the aggregated LHCII as previously observed^8^ and became more pronounced in the dialyzed LHCII (**Fig. 1d, inset**). This red-shifted emission has been described to be due to changes in pigment interaction and/or local hydrophobic environment^12^, and it was suggested to be weak emission caused by Chl-Chl charge transfer states, which occur during a process of ^1^Chl* de-excitation^8^, although the exact mechanism has recently been debated^13,14^. As discussed below, the relative fluorescence yields between each sample observed at room temperature and 77 K appeared to be different (*i.e*., the aggregated LHCII showed a lower yield at 77 K than that measured at room temperature relative to the yield observed in the solubilized LHCII). In addition, when we measured absorption spectra, the dialyzed LHCII showed decreased absorbance at both Soret (400-470 nm) and Qy bands (**Fig. 1e**). Because we used the same Chl concentration of each sample (based on measurement of Chls extracted in acetone), these LHCII samples contain an equal amount of proteins and pigments. In fact, resolubilization of the dialyzed LHCII increased the absorption, which was visibly noticeable (**Fig. 1f**). This change in absorption by aggregation is due to the optical distortion caused by two phenomena: light scattering and inhomogeneous distribution of pigments, the so-called sieve effect^15-17^. Theoretical and experimental studies have shown the complexity of the optical distortion that involves light scattering properties^17-20^. Because LHCII aggregates can scatter a considerable amount of light and also create the sieve situation in which Chls are not distributed homogeneously within the sample^20^, the total probability of photoexcitation per volume becomes smaller than that of solubilized LHCIIs (*i.e*., the incident light is scattered and leaving the sample cuvette before being absorbed by Chls). Similar aggregation-induced changes in optical properties have been observed in non-photosynthetic proteins as well^21,22^. Although we cannot exclude the involvement of other possible mechanisms (*e.g*. Chl concentration quenching^23^), our observations suggest that the aggregation-induced changes in the absorption and light scattering properties of LHCII, independent of the amount of prior light exposure, significantly affects the observed decline of Chl fluorescence intensity.

### The level of aggregation and the amount of light exposure separately affect light-induced fluorescence quenching in LHCII

To investigate how the amount of light exposure impacts on the results shown in Fig. 1, we next observed the kinetics of Chl fluorescence under actinic light (AL) illumination. The solubilized LHCII showed a noticeable Chl fluorescence quenching induced by AL illumination (**Fig. 2a**). This light-induced quenching is considered to be caused by conformational changes within LHCII^24^, which are suggested to promote singlet EET to lutein^6^ (and zeaxanthin^4,5,7^, which is generated from violaxanthin by a violaxanthin de-epoxidase under HL conditions^25^, although zeaxanthin did not exist in any of our LHCII samples isolated after overnight incubation of leaves in the dark). In addition, in isolated LHCII without efficient energy trapping by RCs, the yield of ^3^Chl* becomes significantly increased^3^, and the population of triplet excited state of Car (^3^Car*) generated by triplet EET from ^3^Chl* also becomes significantly higher under HL conditions^26-29^. Despite the relatively long lifetime of ^3^Chl* in a range of tens to hundreds of microseconds^30^, the interaction with Cars, especially within LHCIIs, drastically shortens the lifetime to the subnanosecond order due to efficient triplet EET from ^3^Chl* to a neighboring Car^31-33^. The ultrafast triplet EET in LHCII has been suggested to have evolved to minimize the production of singlet oxygen in oxygenic photosynthetic systems^32,33^. Triplet EET occurs via the Dexter mechanism, and Cars located in L1 and L2 sites (*e.g*. lutein) are considered to be the acceptors of the triplet excitation energy in LHCII^34-36^. This implies that about a third of the total Chls in LHCII are able to undergo de-excitation by the ^3^Car*, the so-called singlet-triplet (S-T) annihilation^28,29^. Thus, it is reasonable to consider that Chl fluorescence quenching in LHCII can occur through both singlet and triplet EET to lutein, especially in isolated conditions under HL illumination.

**Fig. 2.**
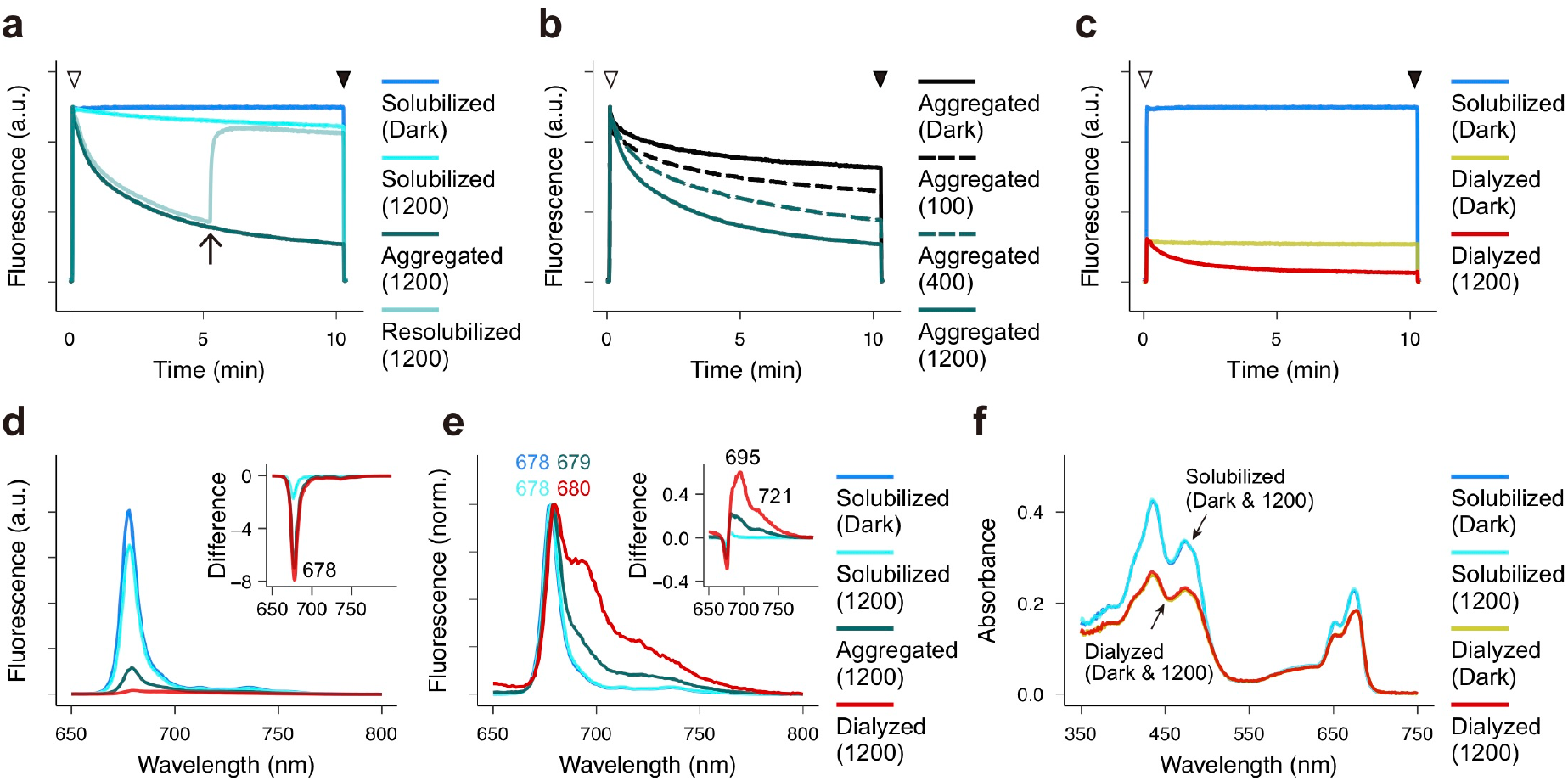
Light-induced fluorescence quenching enhanced by aggregation. **a-c**, Representative kinetics of Chl fluorescence yields observed using different LHCII samples under AL illumination. The preparation of solubilized LHCII, aggregated LHCII, resolubilized LHCII, and dialyzed LHCII was done as described in Fig. 1a. Chl concentration of each LHCII sample was adjusted to 5 µg Chl/mL quantified by extraction in acetone. The light intensity of ML was ∼1 µmol photons m^−2^ s^−1^. An arrow in (**a**) indicates the time point when 0.03% (w/v) α-DDM was added to the aggregated LHCII for resolubilization. Open and closed arrowheads are the points when AL was turned on and off, respectively. The number indicated in sample legend is the light intensity of AL in µmol photons m^−2^ s^−1^. **d**,**e**, Representative Chl fluorescence spectra measured at 77 K normalized at the fluorescence peak (530 nm) of Alexa Fluor 430 (2 nM) added to the sample as an internal standard (**d**) or the main Qy peak (**e**). Excitation wavelength was 440 nm. The insets show the difference spectra from the spectra of the solubilized LHCII (Dark). **f**, Absorbance spectra measured using solubilized LHCII and dialyzed LHCII before (Dark) and after AL illumination at 1200 µmol photons m^−2^ s^−1^ for 10 min. Each sample was adjusted to equal Chl concentration quantified by extraction in acetone. Spectra are shown without normalization.

On the other hand, the extent of the quenching observed in the aggregated LHCII induced by AL illumination was markedly larger than that observed in solubilized LHCII (**Fig. 2a**). Also, the extent of quenching increased with the intensity of AL (**Fig. 2b**). The resolubilization of the aggregated LHCII under AL caused a rapid increase in the fluorescence yield to a level similar to that observed in the solubilized LHCII under the same light intensity (**Fig. 2a**). The dialyzed LHCII also showed additional quenching induced by AL illumination, even though no more aggregation process occurred (**Fig. 2c**). This result differentiates the effect of the aggregation process from that of light illumination, indicating that the light illumination caused the additional quenching within the aggregated structure. We did not observe a distinct difference in the aggregation-induced red-shifted emission (ca. 720 nm) at 77 K between the AL condition and the dark (**Fig. 2d, e** as compared to **Fig. 1c, d**). Also, there was no additional optical distortion induced by HL (**Fig. 2f**). These observations suggest at least two possible explanations: i) the light-induced quenching occurs near the interface between each LHCII in the aggregate; and/or ii) the interaction of pigments at the interface allows an excitation energy migration among LHCIIs in the aggregate, increasing the connectivity to available quenchers. The former possibility can be aligned with the aggregation-induced conformational changes^24^, which allow singlet EET to lutein^6^. The latter possibility is supported by the observations of annihilation processes, which occur significantly more in aggregated LHCIIs than in solubilized LHCIIs^37-40^. Such enhanced annihilation is also observed more in dimeric photosystem II complexes than in the monomeric complexes^41^. The quenching by annihilation processes, especially involving the triplet excited state (*i.e*., S-T and triplet-triplet (T-T) annihilation), may explain the residual quenching after resolubilization, which is caused by the long-lived triplets that remain under HL conditions.

Based on these observations, there are at least three different aspects that need to be taken into account when the level of quenching in LHCII is measured *in vitro*. First, the extent of aggregation needs to be evaluated to determine the level of quenching. Not only does the concentration of detergents affect the extent of aggregation, but the solution pH also greatly affects micelle formation. Thus, the pH-shift treatment from neutral to acidic needs to be carefully analyzed because it would change the level of aggregation. Second, the amount of light exposure needs to be carefully determined. As shown in Fig. 2b, different AL intensities and durations of light exposure generate different levels of quenching. Thus, the level of quenching needs to be determined by comparing before and after a certain amount of light exposure. Third, the involvement of triplet EET in quenching in aggregated LHCII needs to be considered. Previously, observations of triplet EET have been mostly done by ultrafast spectroscopy. However, AL treatments and saturation pulses generally used in modulation fluorometry would be strong enough to generate ^3^Chl*. It is well known that the triplet excited state of Chls and Cars will cause different optical properties^3,34^. In the next section, we took a different approach by using microscopy techniques to examine changes in optical properties of LHCII induced by excess light energy.

### A newly observed radiative process occurs in the green region simultaneously with Chl de-excitation

In relation to the triplet EET described above, we observed a curious emission in the green region from LHCII. We solubilized thylakoid membrane proteins with a mild detergent and separated them by native protein gel electrophoresis. The fluorescence image of the gel indicated that the protein complexes showed emission signals from not only the Chl fluorescence region but also the green region, especially more visible from LHC-containing bands (**Fig. 3a**). Intriguingly, when we illuminated a part of the native gel containing the aggregated LHCIIs for 1 min using a confocal laser scanning microscope, not only did Chl fluorescence yields decrease, but also the intensity of the green emission became stronger than that outside of the illuminated region (**Fig. 3b**). The same green emission was observed in chloroplasts under normal conditions *in vivo* (**Fig. 3c**). When a part of the chloroplast was strongly excited for 0.1 second, the Chl fluorescence yield in the strongly excited region became lowered, while the green emission became stronger in parallel (**Fig. 3d, e; Supplementary Video 1**). It is also evident that the Chl fluorescence yields in the background gradually decreased, but the green emission in the background stayed almost unchanged (**Fig. 3e**). The different kinetics in these regions can be explained by the different yields of ^3^Chl* generated by different amounts of light exposure. The observation laser mainly caused a gradual induction of singlet EET quenching, as observed in the background. On the other hand, the strong excitation laser instantly saturated the RCs and caused an immediate rise of the ^3^Chl* yields in the strongly excited region. The excitation energy of ^3^Chl* cannot be emitted as fluorescence but can be transferred to neighboring Car, generating ^3^Car* and causing Chl fluorescence quenching only in the strongly excited region. The similar kinetics has been observed previously during the triplet EET between ^3^Chl* and ^3^Car*^31-33^.

**Fig. 3.**
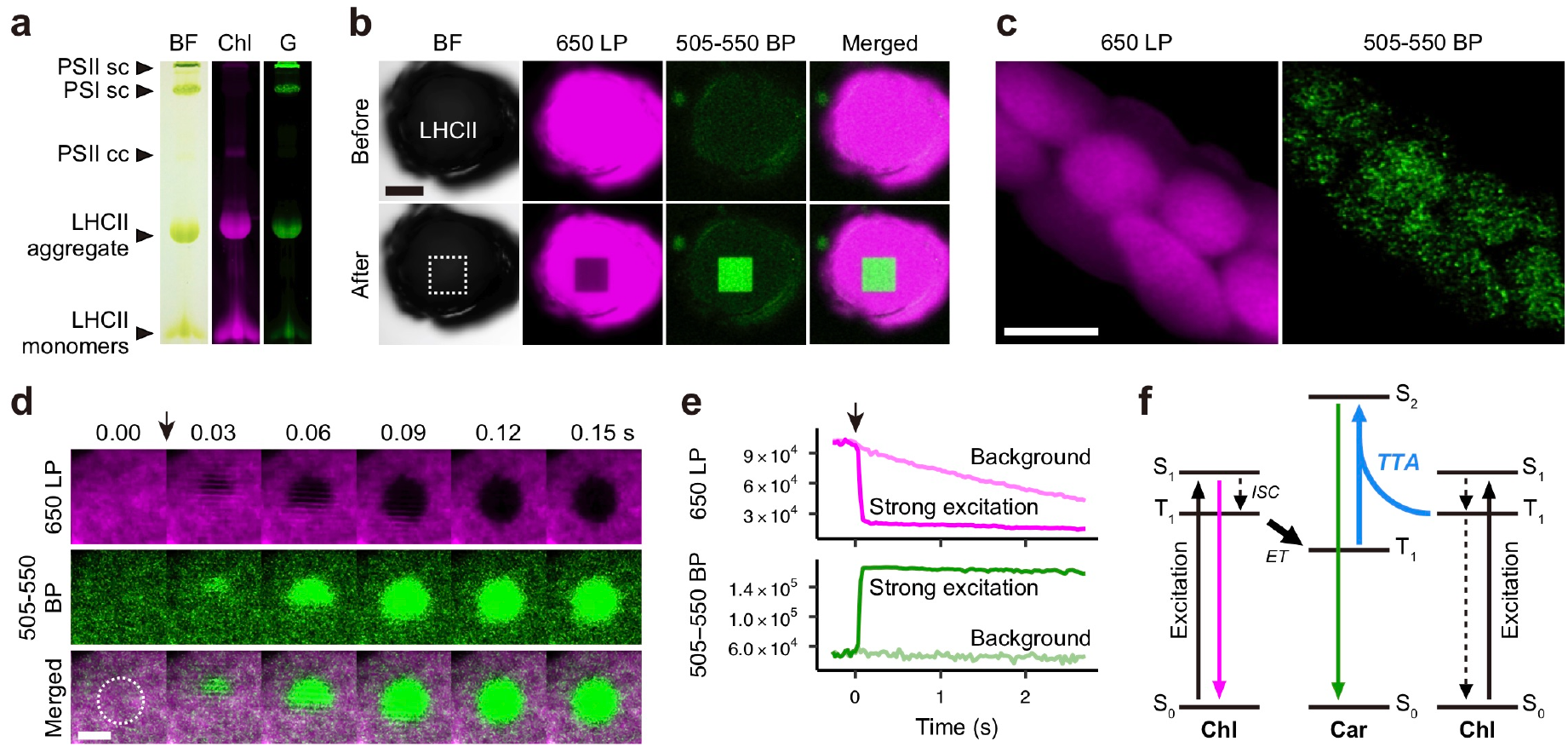
An increased intensity of the green emission caused by excess light energy. **a**, Solubilized thylakoid membrane protein complexes were separated by native gel electrophoresis (see Methods for details). BF, bright-field image of the gel; Chl, the signal observed through a 670 ± 30 nm bandpass (BP) filter; G, the signal observed through a 520 ± 20 nm BP filter. Excitation was 488 nm. **b**, Confocal images of a gel piece from the aggregated LHCII band separated in native gel as shown in (a). After an additional 1-min observation of a square area (indicated by dotted line), Chl fluorescence observed through a 650 nm longpass (LP) filter decreased, and the intensity of the green emission through a 505-550 nm BP filter became stronger than that outside of the square. Scale bar, 100 µm. **c**, Confocal image of chloroplasts in a protonema cell of *Physcomitrella patens*, showing Chl fluorescence (650 LP) and the green emission (505-550 BP). Excitation was 488 nm. Scale bar, 10 µm. **d**, Real-time imaging of local, strong excitation (indicated as dotted circle) in a chloroplast showing a simultaneous decrease and increase in the intensity of Chl fluorescence (650 LP) and the green emission (505-550 BP), respectively. The arrow indicates the starting point of local, strong excitation (for 0.1 s) using a 405 nm laser. Excitation for real-time observation was 488 nm. Scale bar, 5 µm. **e**, Traces of the averaged intensity of emission observed in the strongly excited region and background as in (d). The arrow indicates the starting point of local, strong excitation (for 0.1 s). See also Supplementary Video 1. **f**, An energy-level diagram for the green emission caused by excess light energy. Photoexcitation prepares the singlet excited state of Chl (S_1_ Chl), which evolves into the triplet excited state (T_1_ Chl) through intersystem crossing (*ISC*). Triplet Chl undergoes Dexter-type energy transfer (*ET*) and eventually generates the triplet excited state of Car (T_1_ Car). Under excess light conditions, T-T annihilation (*TTA*) takes place between T_1_ Chl and T_1_ Car and yields the singlet excited state of Car (S_2_ Car) which can show green emission. It should be noted that T_1_ Car can also induce S-T annihilation which quenches the red emission from S_1_ Chl (not shown in the diagram for simplicity).

The question is whether the coinciding rise in the green emission is related to the ^3^Car* de-excitation or not. It is generally considered that carotenoids are non-emissive; however, a weak fluorescence was reported previously^42^. Especially when Cars contain more than around eight conjugated double bonds, they exhibit emission from the second electronically excited singlet state (S_2_)^43^. The spectral deconvolution of LHCII indicates that the S_2_ energies of lutein at L1 and L2 sites and violaxanthin are 20250, 20050, and 20200 cm^−1^, respectively^44^. The excited triplet state (T_1_) energy of β-carotene is estimated to be 8000–8900 cm^−1 45,46^. Given the T_1_ energies of Chl *a* and *b* (10700 and 11200 cm^−1^, respectively)^47^, the upconverted energies through T-T annihilation are similar to the S_2_ energies of carotenoids (18700–20100 cm^−1^, **Fig. 3f**). Moreover, it has been shown that T-T annihilation processes can generate photoluminescence^48^. It should also be noted that S-T annihilation between the lowest excited singlet state (S_1_) of Chl and T_1_ Car can also generate a higher excited triplet state (T_n_) of Car, which has a higher energy level than that of S_2_ Car. The yield of intersystem crossing of Car is typically low; however, there is a mixing between excited states of Chl and Car in the plant LHCII^49,50^ which can alter the strength of spin-orbit coupling and emission intensity. Although further investigation is certainly needed for understanding the relationship between the green emission dynamics and S-T and/or T-T annihilation processes, these results demonstrate that another radiative de-excitation pathway exists in isolated LHCII under HL conditions.

### Aggregated LHCIIs in liposome environments show unique optical properties

To further investigate changes in optical properties of LHCII, we next used LHCII incorporated in liposomes, which provides the opportunity to study fluorescence quenching mechanisms under more native conditions^51,52^. We prepared LHCII-containing liposomes (proteoliposomes, or PLs) by detergent-mediated reconstitution (see Methods for details). We removed by-products (*e.g*. smaller or larger PLs, empty liposomes, etc.) by sucrose gradient centrifugation and used only the PLs from the most abundant single band in the gradient (**Supplementary Fig. 1**). We used the same amount of LHCII (100 µg Chl) incorporated in either 1.5 or 0.5 mg of liposomes to generate LHCII-PLs with low or high densities of LHCII per liposome, respectively, whose differences were visible by sucrose gradient (**Supplementary Fig. 1d, e)**.

We examined Chl fluorescence properties of these two different LHCII-PLs in the same way as above. The fluorescence yields observed in these two PLs showed about half of the yield observed in the solubilized LHCII (**Fig. 4a, b**). The high-density PL showed a slightly lower yield than the low-density PL. Upon AL illumination, similar Chl fluorescence quenching was observed in both PLs, indicating that the light-induced fluorescence quenching of LHCII still occurs in lipid membrane conditions. Interestingly, the red-shifted fluorescence emission shoulder measured at 77 K of the low-density PL was actually stronger than that of the solubilized LHCII (**Fig. 4c**), which was not observed in the high-density PL (**Fig. 4d**) or other LHCII samples (**Figs. 1c, 2d**). This could indicate a weakly aggregated state in liposome conditions. Similar to the other LHCII conditions, the fluorescence yield of the shoulder became smaller under AL illumination, indicating that the intensity of total emission actually becomes smaller by the light-induced quenching. The fluorescence spectral shape was similar between the two PLs, although there was a slight shift of the peak between the two (**Fig. 4e, f**), which suggests different levels of aggregation occurring between the two PLs.

**Fig. 4.**
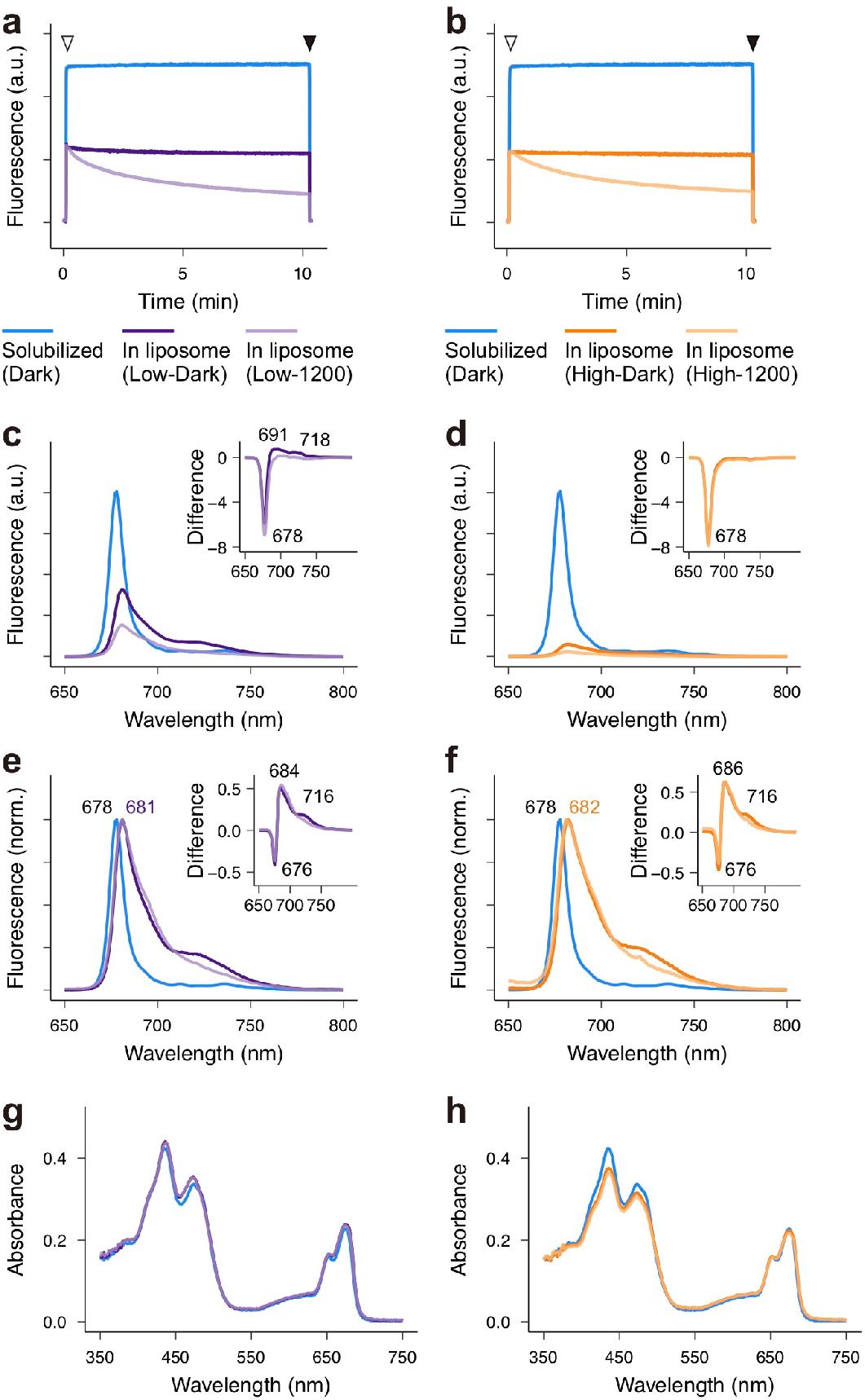
Unique optical properties of LHCII in liposomes. Two different LHCII-PLs containing low (**a**,**c**,**e**,**g**) and high (**b**,**d**,**f**,**h**) densities of LHCII per liposome were prepared (see Methods for details; also see Supplementary Fig. 1). **a**,**b**, Representative kinetics of Chl fluorescence yields observed using the two LHCII-PLs. The preparation of solubilized LHCII was done as described in Fig. 1a. The light intensity of ML is ∼1 µmol photons m^−2^ s^−1^. Open and closed arrowheads are the points when AL (at 1200 µmol photons m^−2^ s^−1^) was turned on and off, respectively. **c**-**f**, Representative Chl fluorescence spectra measured at 77 K normalized at the fluorescence peak (530 nm) of Alexa Fluor 430 (2 nM) added to the sample as an internal standard (**c**,**d**) or the main Qy peak (**e**,**f**). Excitation wavelength was 440 nm. The insets show the difference spectra from the spectra of the solubilized LHCII (Dark). **g**,**h**, Absorbance spectra measured using the solubilized LHCII and the LHCII in liposomes before (Dark) and after AL illumination (at 1200 µmol photons m^−2^ s^−1^) for 10 min. Spectra are shown without normalization. Chl concentration of each LHCII sample was adjusted to 5 µg Chl/mL quantified by extraction in acetone. Line colors were the same as in (**a**,**b**).

As shown above, Chl fluorescence yields observed in the low- and high-density LHCII-PLs were only slightly different from each other at room temperature (**Fig. 4a, b**). However, their relative fluorescence yields observed at 77 K showed a distinct difference—the yield for the high-density PL was about 80% lower than that of the low-density PL (**Fig. 4c, d**). This discrepancy is likely influenced by different levels of light scattering caused by different levels of aggregation between the two PLs. Another critical difference was the optical setup between modulation fluorometry and spectroscopy at 77 K. The signals observed by using a modulation fluorometer at room temperature were directly collected from the sample cuvette through the light guide to the photodiode unit. On the other hand, the fluorescence measurement at 77 K was done using a spectrophotometer, in which the cryogenic sample dewar was located distantly from the photomultiplier tube with a 2-nm slit. Because of these differences in optics and the sample-detector distance, the signal detection appeared to be influenced by different levels of light scattering in the samples. As mentioned earlier, the same situation was observed in Fig. 1. The observed discrepancy highlights the following questions: i) how specific and complementary are the signals observed by using these two methods and ii) how much does light scattering impact fluorescence measurements in different experimental setups. Because of the complexity in determining the contributions of absorption and light scattering^15,17-20^ which influence fluorescence and the quenching, further investigation is required to address these issues.

It should be noted that the absorbance spectra in this study were measured by using an integrating sphere, in which the sample was located in front of the entrance port. When the amount of backscatter (reflectance) is negligible, the total transmitted light collected within the sphere will provide true absorption spectra. However, the true absorption spectrum of dialyzed LHCII was not similar to that of the solubilized LHCII (**Figs. 1e, 2f**), whereas that of the LHCII-PLs was similar to that of solubilized LHCII (**Fig. 4g, h**). This difference may indicate that the contributions of the sieve effect caused by LHCII aggregation are minimized by being situated in the lipid environments of PLs. Thus, it is very likely that the aggregated LHCII *in vitro* and LHCII in lipid environments (*e.g*. PLs or thylakoid membranes) have different optical properties, which also affect their fluorescence properties.

### Variable optical properties of LHCII impact observed quenching mechanisms

In this study, we investigated fluorescence and absorption properties of LHCII in different physical conditions based on equal Chl concentrations. We conducted this study because experimental light illumination (*e.g*. AL or laser) caused changes in fluorescence lifetime of LHCII samples *in vitro* as initially observed by Jennings *et al*.^53^. For example, when we used LHCII-PLs for time-correlated single-photon counting measurements, the fluorescence lifetime became shorter from ∼1.7 to ∼1.5 ns during the three consecutive measurements (a few seconds for each measurement). When we illuminated the LHCII-PLs with AL at ∼850 µmol photons m^−2^ s^−1^ for 10 min, the lifetime became distinctly shorter to ∼0.6 ns. This phenomenon is quite problematic because light-induced quenching will be induced differently depending on the amount of light exposure (*e.g*. Fig. 2b), which would be difficult to standardize among different experimental setups in different labs. Therefore, it will be essential to specify the amount of light exposure (*i.e*., wavelength, intensity, and duration) when investigating the level of quenching, especially with *in vitro* samples.

We also revisited light scattering and the sieve effect induced upon LHCII aggregation. Theoretical and experimental techniques can correct the distorted absorption caused by both light scattering and the sieve effect^17-20^. Even so, the correction cannot change the fact that these two phenomena affect how photons interact with LHCIIs and change the amount of light absorption per Chl molecule. To take these two phenomena into consideration, we conducted this study based on equal Chl concentrations. Because of the variable optical properties generated by altered physical conditions, Chl fluorescence yields become different regardless of environmental light conditions (**Fig. 1**). Therefore, it is important to consider that the optical properties (*e.g*. absorption) in the samples can vary during measurements, which could affect fluorescence intensity. The physical nature of LHCII aggregation could influence light harvesting in two contradictory ways *in vivo*: i) lowering light absorption as observed in this study; and ii) increasing the opportunities for scattered light to be absorbed by other LHCs in different membrane regions, similar to the detour effect described in leaf tissue optics^16^. Although numerous studies have shown that light scattering occurs differently in thylakoid membranes under different light conditions^54^, it is still largely unknown how the scattered light influences the total light usage within a chloroplast.

Table 1 summarizes possible mechanisms that decrease Chl fluorescence in different LHCII conditions. Although we cannot simply apply *in vitro* results to *in vivo* situations, we would like to consider the possible situations based on this study. It is evident that the solubilized LHCII shows the highest fluorescence yield among other physical environments examined here. Thus, the extent of quenching observed in the solubilized LHCII is smaller than that of the other LHCII samples *in vitro*. Regardless of environmental light conditions, fluorescence intensities become lower upon aggregation of LHCII due to the decreased level of absorption caused by light scattering and the sieve effect. Although it appears to be fluorescence quenching, this is not related to Chl de-excitation mechanisms but simply results from altered physical conditions of LHCII. Under HL, quenching mechanisms are activated through singlet and triplet EET to other pigments (*e.g*. lutein). Singlet EET has a faster rate constant than that of triplet EET, so the contribution of the quenching via singlet EET will be initially high. On the other hand, under a prolonged HL exposure (as frequently used in modulation fluorometry), the contribution of the quenching of ^3^Car* generated by triplet EET increases. Moreover, we found an increase in the green emission upon Chl fluorescence quenching under HL, which might be related to S-T and/or T-T annihilation processes^43,48-50^ and will need to be taken into consideration for understanding the total energy budget.

**Table 1.** Possible mechanisms that decrease Chl fluorescence in different LHCII conditions

| **LHCII conditions** | **Light scattering (LS) & sieve effect (SE)** | **Quenching via $^1\text{Chl}^*$ de-excitation** | **Quenching via triplet EET ($^3\text{Chl}^* - ^3\text{Car}^*$)** |
| --- | --- | --- | --- |
| Solubilized LHCII | Both LS & SE negligible | Small (*i.e.* high fluorescence) | Only under HL |
| Aggregated/dialyzed LHCII | Both LS & SE increased, Abs. decreased | Active | Enhanced under HL and involved with annihilation processes |
| LHCII in liposome | More LS from liposomes and from LHCII aggregates, but less SE | Active | Enhanced under HL and involved with annihilation processes |
| *In vivo* | Multiple effects by cellular & organellar matrices | Additional regulation by PsbS and enhanced by zeaxanthin | Increased when open RCs become unavailable/inaccessible by chemical inhibitors or mutations |

As previously shown^51,52^, LHCII in liposome environments tends to form aggregates, which most likely represent native conditions in thylakoid membranes. The different concentration of LHCII per liposome still generates different levels of light scattering, which appears to affect fluorescence intensities depending on the measurement setups (**Fig. 4**). The light-induced quenching occurs similarly in LHCII-PLs, suggesting that evaluating fluorescence properties of LHCII-PLs also needs to be standardized based on the amount of light exposure. This observation also suggests that the light-induced quenching involving the triplet EET could be enhanced even *in vivo* when open RCs are not available under HL conditions and also the situation in which the number of RCs per LHCII is reduced by mutations or chemical treatments. Obviously, it is more complicated to dissect the quenching properties *in vivo* because of the presence of monomeric LHCII^55^, xanthophyll cycle^25^, and photosystem II subunit S (PsbS)^56^. Nevertheless, it is more essential to consider collectively how different components have impact on overall quenching mechanisms rather than to focus exclusively on a specific mechanism, especially when we investigate the process of NPQ induction *in vivo*. Furthermore, our study emphasizes that multiple processes that affect Chl fluorescence yields can emerge as the level of protein-protein interactions and the amount of light exposure increase. To dissect how these multiple processes progressively arise under excess light conditions, it is necessary to take genetic approaches (such as the mutants lacking monomeric LHCII^55^ and trimeric LHCII^57^) and consider the spatial and temporal correlation between the different components that contribute to overall quenching mechanisms.

## Methods

### Isolation of LHCII

We obtained spinach leaves from a local store a day before LHCII isolation, and leaves were kept moist with wet paper towels and incubated at 4 °C in the dark overnight. Isolation of LHCII was done as described previously^2^. Deveined leaves were homogenized in 25 mM Tricine-KOH (pH 7.8), 400 mM NaCl, 2 mM MgCl_2_, 0.2 mM benzamidine, and 1 mM ε-aminocaproic acid at 4 °C using a Waring blender for 30 s with max speed. The homogenate was filtered through 4 layers of Miracloth, and the filtrate was centrifuged at 27,000 × g for 10 min at 4 °C. The pellet was resuspended in 25 mM Tricine-KOH (pH 7.8), 150 mM NaCl, 5 mM MgCl_2_, 0.2 mM benzamidine, and 1 mM ε-aminocaproic acid. The suspension was loaded on sucrose cushion containing 1.3 M sucrose with 25 mM Tricine-KOH (pH 7.8), 15 mM NaCl, and 5 mM MgCl_2_, which was overlaid on 1.8 M sucrose with 25 mM Tricine-KOH (pH 7.8), 15 mM NaCl, and 5 mM MgCl_2_, and centrifuged at 131,500 × g for 30 min at 4 °C using a SW 32 Ti rotor (Beckman Coulter). Thylakoid membranes sedimented in 1.3 M sucrose cushion were collected and washed with 25 mM Tricine-KOH (pH 7.8), 15 mM NaCl, and 5 mM MgCl_2_, and centrifuged at 27,000 × g for 15 min at 4 °C. The pellet was resuspended in 25 mM Tricine-KOH (pH 7.8), 0.4 M sucrose, 15 mM NaCl, and 5 mM MgCl_2_, and centrifuged at 27,000 × g for 10 min at 4 °C. The pellet was resuspended and used as purified thylakoid membranes. The purified thylakoid membranes were resuspended in 25 mM HEPES-NaOH (pH 7.8) and centrifuged at 15,300 × g for 10 min at 4 °C. The pellet was resuspended in 25 mM HEPES-NaOH (pH 7.8) at 2.0 mg Chl/mL and solubilized with 4% (w/v) n-dodecyl-α-D-maltoside (α-DDM as previously abbreviated as α-DM; Anatrace) for 30 min with gentle agitation on ice. The unsolubilized membranes were removed by centrifuging at 21,000 × g for 5 min at 4 °C. The supernatant was filtered through 0.22 μm filter using Durapore Ultrafree filters centrifuged at 10,000 × g for 3 min at 4 °C. The 200 μL of filtered solubilized fraction was used for gel filtration chromatography using the ÄKTAmicro chromatography system with a Superdex 200 Increase 10/300 GL column (GE Healthcare) equilibrated with 25 mM HEPES-NaOH (pH 7.8) and 0.03% (w/v) α-DDM at room temperature. The flow rate was 0.9 mL/min. The proteins were detected at 280 nm absorbance. The fraction separated from 10.0 to 10.3 mL contained trimeric LHCII proteins. Several runs (typically 3-5 times) were performed consecutively to obtain a sufficient amount of sample.

### Modulation fluorometry

Chl concentration of each type of LHCII sample was pre-determined by extracting Chl in 80% acetone as described previously^58^. The Chl concentration of each sample was adjusted to 5.0 µg Chl/mL in 25 mM HEPES-NaOH (pH 7.8) with 0.4 M sucrose. Chl fluorescence kinetics was measured using Dual-PAM-100 (Walz). The intensity of ML was set to 7 (∼1 µmol photons m^−2^ s^−1^). For Fig. 1b, the intensity setting 2 (∼0.5 µmol photons m^−2^ s^−1^) and 12 (∼5 µmol photons m^−2^ s^−1^) were also used. The intensity of AL (red light) was set to 15 (∼1200 µmol photons m^−2^ s^−1^) with Emitter DUAL-E. For Fig. 2b, the intensity setting 5 (∼100 µmol photons m^−2^ s^−1^) and 10 (∼400 µmol photons m^−2^ s^−1^) were also used. Detector DUAL-DR was used to monitor Chl fluorescence emission (> 700 nm). The sample was continuously stirred using a magnetic stir bar.

### Fluorescence spectroscopy at 77 K

Chl concentration of each type of sample was pre-determined by extracted Chl in 80% acetone as described previously^58^. The Chl concentration of each sample was adjusted to 5 µg Chl/mL in 25 mM HEPES-NaOH (pH 7.8) with 0.4 M sucrose. For absolute quantitative comparison, 2 nM Alexa Fluor 430 (Thermo Fisher Scientific) was added to each sample and mixed with a magnetic stir bar for 15 s. The sample was placed in a glass tube and frozen in liquid nitrogen for 5 min. Fluorescence emission was recorded at 77 K using FluoroMax-4 spectrophotometer (Horiba Scientific). Excitation wavelength was 440 nm with a 2-nm slit size. Emission wavelength measured was from 500 to 800 nm with a 2-nm slit size. Fluorescence emission for each sample was recorded consecutively three times to obtain average spectra.

### Absorption spectroscopy

Chl concentration of each type of sample was pre-determined by using the extracted Chl in 80% acetone as described previously^58^. For measurements using an integrating sphere, the Chl concentration of each sample was adjusted to 50 µg Chl/mL in 25 mM HEPES-NaOH (pH 7.8) with 0.4 M sucrose. Total transmittance from 350 to 750 nm (1 nm step size) was recorded using a quartz cuvette with 1-mm pathlength located in front of the entrance port of the integrating sphere installed in a LAMBDA 950 UV/Vis spectrophotometer (PerkinElmer). A blank was recorded in each measurement using 25 mM HEPES-NaOH (pH 7.8) with 0.4 M sucrose.

### Native protein gel electrophoresis

Isolated thylakoid membranes were solubilized with 2% (w/v) α-DDM for 30 min on ice. Insoluble fractions were removed by centrifugation at 21,000 × g for 5 min at 4 °C. The solubilized fraction was subjected to clear native polyacrylamide gel electrophoresis as described previously^59^. The sample was applied to a 5-13% (w/v) acrylamide gradient gel with 25 mM imidazole-HCl (pH 7.0) as anode buffer and 50 mM Tricine-HCl (pH 7.0), 7.5 mM imidazole, 0.02% α-DDM, and 0.05% deoxycholate as cathode buffer. Fluorescence images of the gel were acquired using a Typhoon 9400 imager (GE Healthcare) using excitation at 488 nm. The signals from the green region and Chl fluorescence were measured using a 520 ± 20 nm bandpass filter and a 670 ± 30 nm bandpass filter, respectively.

### Confocal laser scanning microscopy

The small piece of the native protein gel containing aggregated LHCII was observed by using a Zeiss LSM510 confocal laser scanning microscope with a Plan Apochromat 63×/1.4 NA oil objective lens. The excitation was at 488 nm, and a 505-550 nm bandpass filter and a 650 nm longpass filter were used to observe the signals from the green region and Chl fluorescence, respectively. After an initial scan, a square region within the gel was selected and continuously scanned for 1 min (**Fig. 3b**). After that, the original image area was restored and scanned again to see the difference in the signal intensity between the green region and Chl fluorescence. Using the same microscopy condition, the chloroplasts in *Physcomitrella patens* protonema tissue were observed as previously described^60^. With a larger size of chloroplast induced by ampicillin as described previously^60^, the simultaneous strong excitation and observation was done using an Olympus FV1000 confocal laser-scanning microscope. The excitation for observation was at 488 nm, and a 505-550 nm bandpass filter and a 650 nm longpass filter were used to observe the signals from the green region and Chl fluorescence, respectively. Scan speed was at 33 frames per second. Strong excitation was done for 0.1 s using a 405 nm laser. Image analysis was done using ImageJ software (US National Institutes of Health).

### Preparation of LHCII proteoliposomes

To prepare liposomes, we used 1,2-dioleoyl-sn-glycero-3-phosphocholine (DOPC: Avanti Polar Lipids) and 1,2-dioleoyl-3-trimethylammonium-propane (DOTAP: Avanti Polar Lipids). The dried DOPC and DOTAP were resuspended in chloroform:methanol (2:1) at the ratio of DOPC:DOTAP = 80:20 (mol%). The lipid mixture was dried for 30 min under stream of nitrogen gas at 40 °C. The dried lipids were resuspended in 25 mM HEPES-NaOH (pH 7.8) and 0.03% (w/v) α-DDM at 40 °C and vortex vigorously for 3 times of 30 s. The lipid solution was warmed back for 30 s at 40 °C in each interval. The resuspended crude liposomes were extruded through a filter with 100-nm pore size for 21 times using LiposoFast. The extruded liposomes were stored at 4 °C until use. Incorporation of solubilized LHCII into liposomes was done by a detergent-mediated dialysis method. The concentration of LHCII in the liposome-LHCII mixture was adjusted to 100 µg Chl/mL. The concentration of liposomes in the liposome-LHCII mixture was adjusted to 1.5 or 0.5 mg/mL to generate the lighter or denser PLs, respectively. The volume of liposome-LHCII mixture was adjusted to 1 mL using 25 mM HEPES-NaOH (pH 7.8) and 0.03% (w/v) α-DDM and dialyzed in 25 mM HEPES-NaOH (pH 7.8) using Slide-A-Lyzer 20K MWCO G2 Dialysis Cassettes (Thermo Fisher). Dialysis was performed at 4 °C, stirred with a magnetic stir bar in the dark. Dialysis buffer was refreshed as follows: 1 L/sample for 1 h, fresh 1 L/sample for 15 h, fresh 1 L/sample for 4 h, and fresh 1 L/sample for 4 h. The dialyzed samples were loaded on sucrose gradient (each concentration overlaid with the denser one: 2 mL of 0.1, 0.4, 0.7, 1.0, and 1.3 M sucrose with 25 mM HEPES-NaOH, pH 7.8) in an ultracentrifuge tube. The sucrose gradient was centrifuged at 154,300 × g for 15 h at 4 °C using a SW 41 Ti rotor (Beckman Coulter). The LHCII proteoliposomes separated from empty liposomes were collected dropwise from the bottom of the tube and used for analysis.

### Fluorescence lifetime measurements

Chl fluorescence lifetimes of LHCII-PLs were measured using a home-built fluorescence lifetime measurement apparatus as described previously^61^. The mode locked femtosecond laser at 840 nm was generated by a Ti:Sapphire oscillator (Coherent Mira 900) at 76 MHz repetition rate. The resulting light was frequency-doubled to 420 nm with a β-barium borate crystal to excite Chls. The two light paths were directed: one for the SYNC pulse for the TCSPC card (Becker-Hickl SPC-630 and SPC-850) and the other for sample excitation. The resulting instrument response function had a full width at half maximum of 40–50 ps. The Chl fluorescence from the sample was directed through a polarizer, followed by a 600-nm longpass filter, to a monochromator (Horiba Jobin-Ivon H-20) with the detection wavelength at 680 nm. The detection system was composed of an MCP/PMT detector (Hamamatsu R3809U), electrically cooled to -30 °C, and connected to a DCC-100 detector control card (Becker-Hickl). A computer with a TCSPC module (Becker and Hickl GmbH, SPC-600) was used for data collection and processing.

## Data availability

All data are available from the corresponding author upon request.

## Acknowledgements

We are grateful to Collin Steen and Dhruv Patel for critical reading of the manuscript. We thank Jacob Jonsson for technical assistance with absorption spectroscopy with an integrating sphere. Y.Y. appreciates the support of the Japan Society for the Promotion of Science (JSPS) Postdoctoral Fellowship for Research Abroad. This work was supported by the US Department of Energy, Office of Science, Basic Energy Sciences, Chemical Sciences, Geosciences, and Biosciences Division under Field Work Proposal 449B (G.R.F. and K.K.N.). K.K.N. is an investigator of the Howard Hughes Medical Institute.

## Author contributions

M.I., G.R.F., and K.K.N. conceived the study. M.I., J-J.C., and S.P. conducted research. M.I., J-J.C., S.P., Y.Y., E.M.S. analyzed data. D.A.F., G.R.F., and K.K.N. provided resources and supervision. M.I. wrote the paper. All authors discussed the results and commented on the manuscript.

## Competing interests

The authors declare no competing interests.

**Correspondence** should be addressed to M.I.

**Supplementary Fig. 1.**
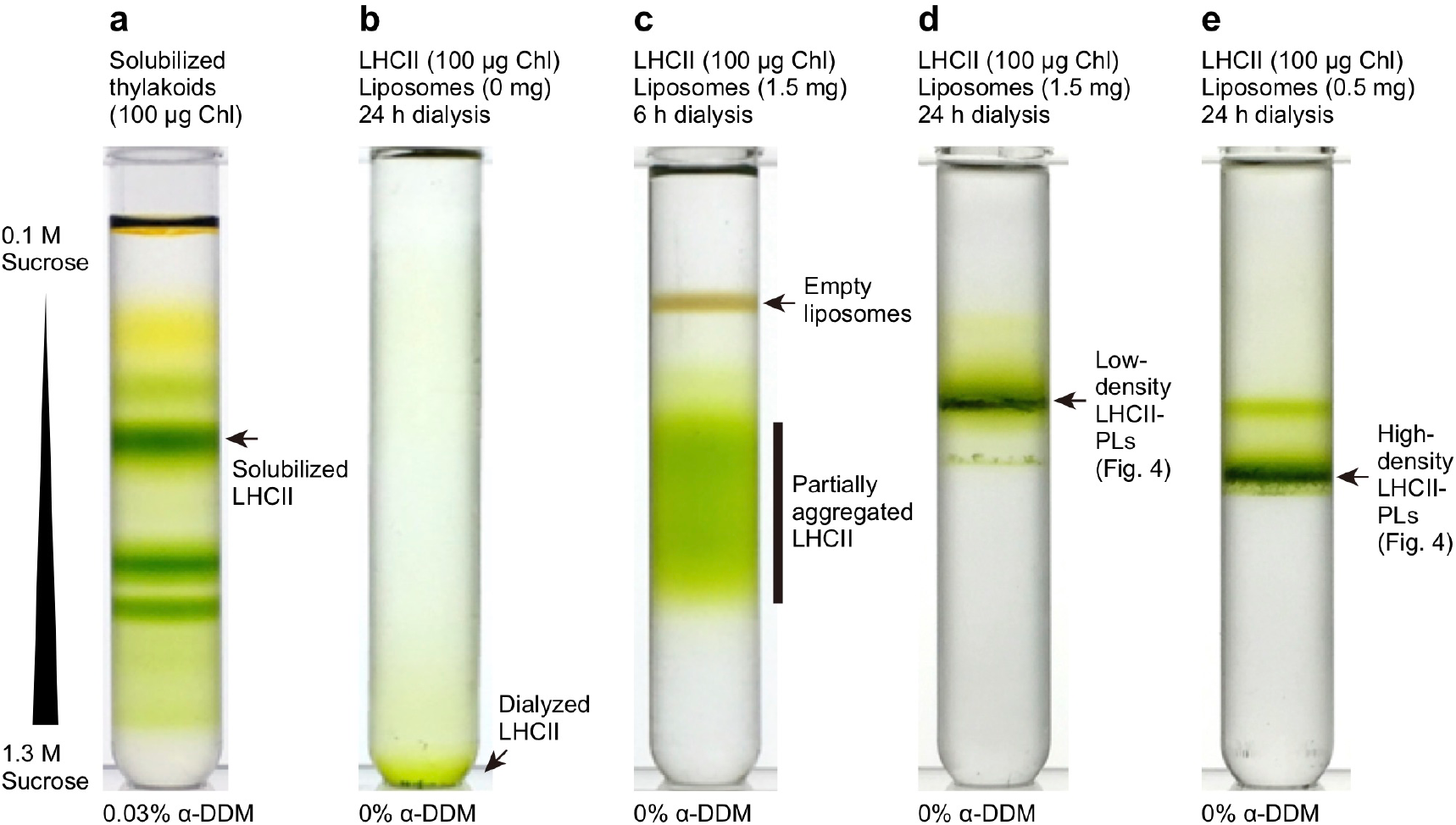
Preparation of LHCII-PLs. We used detergent-mediated reconstitution to prepare LHCII-containing PLs (see Methods for details). We used sucrose gradient to separate and purify LHCII-PLs (see Methods for details). **a**, As a control, solubilized thylakoids were separated by sucrose gradients, which contained 0.03% α-DDM. Solubilized LHCIIs were located at the arrow. **b**, Solubilized LHCIIs were dialyzed for 24 h to remove α-DDM completely and separated by sucrose gradients, which did not contain 0.03% α-DDM. Because of the lack of detergent, dialyzed LHCIIs were sedimented at the bottom of the gradient. **c**, Solubilized LHCIIs and liposomes were mixed and subjected to dialysis for 6 h. Because of the incomplete dialysis, the incorporation of LHCIIs into liposomes did not occur so that the empty liposomes and partially aggregated LHCIIs were separated in the gradients. **d**,**e**, Solubilized LHCIIs and 1.5 mg (**d**) and 0.5 mg (**e**) of liposomes were mixed and dialyzed for 24 h. Sucrose gradients separated the low and high-density LHCII-PLs. Because of the difference in densities, the two PLs were sedimented at different locations in the gradients. The detergent concentration indicated below each figure is the concentration in the sucrose gradients.

**Supplementary Video 1.**
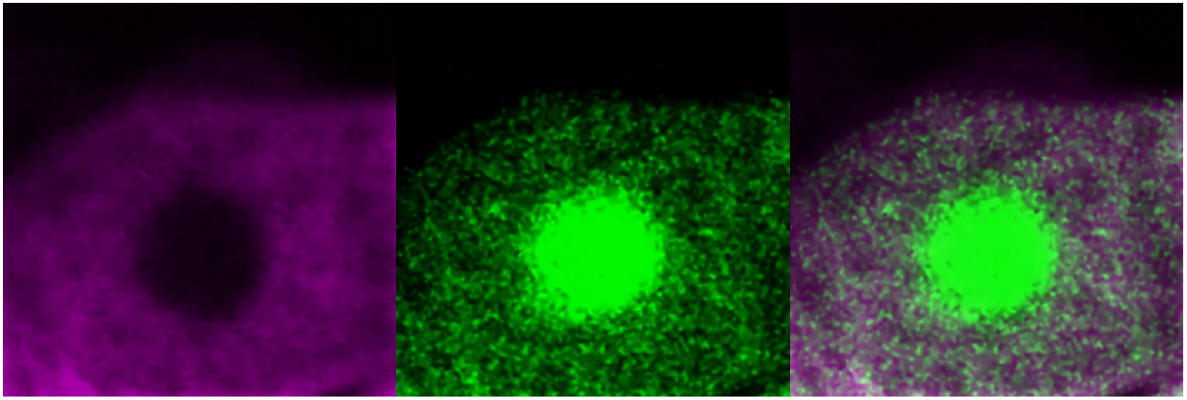
Real-time imaging of local, strong excitation in a chloroplast (Fig. 3d). Confocal images of chloroplasts in a protonema cell of *Physcomitrella patens*, showing Chl fluorescence (left, 650 LP), the green emission (middle, 505-550 BP), and merged image (right). Excitation for real-time observation was 488 nm. The local, strong excitation was done for 0.1 s with a 405 laser. See Methods for details.

